# Disentangling the role of surface topography and intrinsic wettability in the prey capture mechanism of *Nepenthes* pitcher plants

**DOI:** 10.1101/2020.10.09.332916

**Authors:** David Labonte, Adam Robinson, Ulrike Bauer, Walter Federle

## Abstract

*Nepenthes* pitcher plants capture prey with leaves specialised as pitfall traps. Insects are trapped when they ‘aquaplane’ on the pitcher rim (peristome), a surface structured with macroscopic and microscopic radial ridges. What is the functional significance of this hierarchical surface topography? Here, we use insect pad friction measurements, photolithography, wetting experiments and physical modelling to demonstrate that the ridges enhance the traps’ efficacy by satisfying two functional demands on prey capture: Macroscopic ridges restrict lateral but enhance radial spreading of water, thereby creating continuous slippery tracks which facilitate prey capture when little water is present. Microscopic ridges, in turn, ensure that the water film between insect pad and peristome remains stable, causing insects to aquaplane. In combination, the hierarchical ridge structure hence renders the peristome wettable, and water films continuous, so avoiding the need for a strongly hydrophilic surface chemistry, which would compromise resistance to desiccation and attract detrimental contamination.

## Introduction

Many plant surfaces interact with water to fulfil biologically important tasks. For example, plants famously use surfaces with remarkable wetting properties to float on water [1], to attain ‘self-cleaning’ properties [2], or for directional transport of water [3–5]. These wetting properties are usually achieved through a combination of intricate surface topographies and the specific surface chemistry of the plant cuticle [6, 7], which covers most primary plant surfaces, and serves as a water-proofing layer allowing plants to thrive in dry environments [8–11]. Because of this functional role, the plant cuticle is usually hydrophobic; plant surface patterned with microscopic surface topographies are thus often highly water-repellent [12–14]. There are however some notable, albeit less well-studied, examples of wettable plant surfaces [15, 16].

A remarkable example of an extremely wettable plant surface is found in carnivorous *Nepenthes* pitcher plants, where a specialised superhydrophilic surface on the pitcher rim (peristome) is essential for prey capture, as it stabilises thin water films on which insects aquaplane [17, 18, see Fig. 1 A]. This slippery surface has recently inspired the development of ‘omni-phobic’ synthetic coatings to which virtually nothing sticks [19, 20]. However, and in sharp contrast to the synthetic surfaces it inspired, the peristome does not trap a wetting liquid with low surface tension (typically perfluorinated lubricants), but a polar liquid with high surface tension (water). How exactly thin layers of water can be stabilised on the pitcher peristome without a strongly hydrophilic surface chemistry that would compromise the water-proofing function of the cuticle remains an open question.

**Figure 1.**
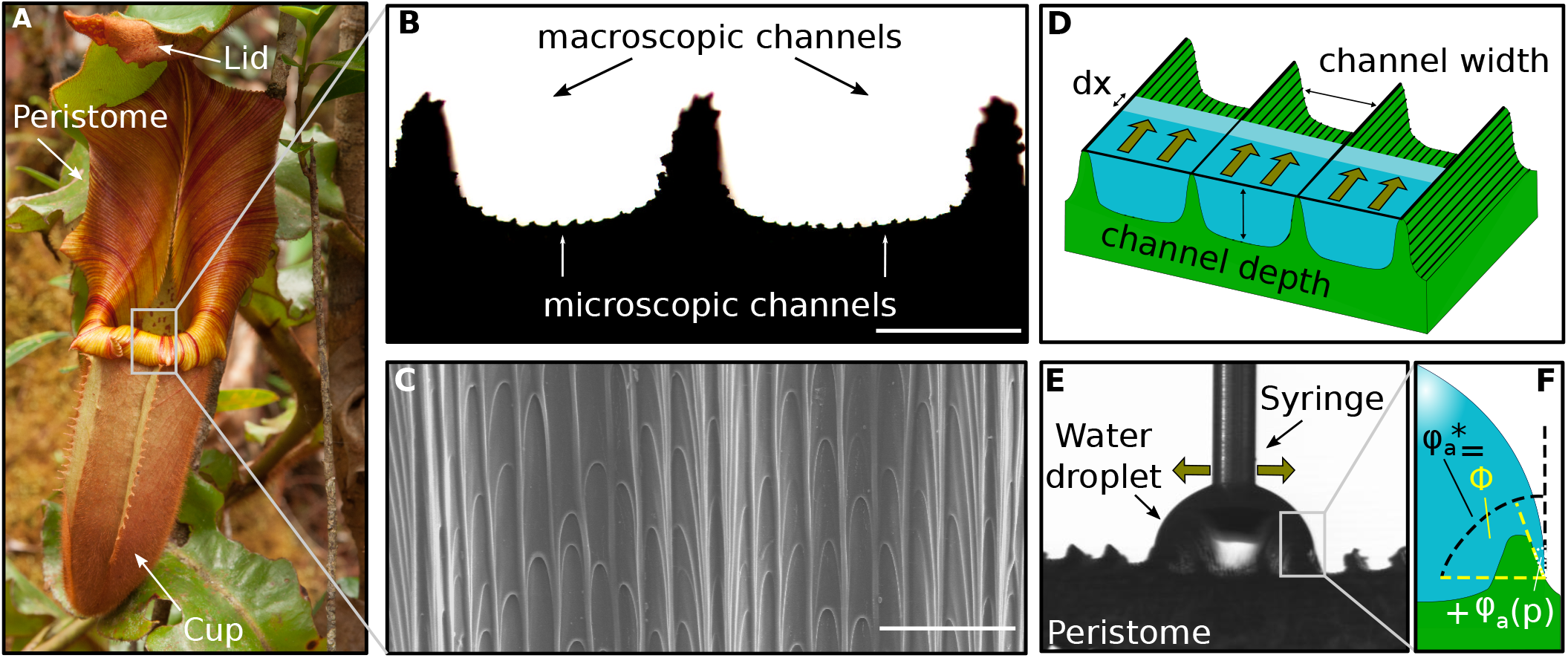
(A) Pitcher plants (here *Nepenthes veitchii*) capture insects by means of a passive pitfall trap which relies on a specialised slippery surface – the peristome. (B & C) Most pitcher peristomes are covered by characteristic channel-like patterns at two different length-scales (macroscopic and microscopic channels, highlighted by black and white arrows, respectively). (B) shows a light microscopy image of a cross-section, whereas (C) shows a scanning electron micrograph of a top-down view (both scale bars 100 μm). (D) Macroscopic channels render spreading of water along the channels energetically favourable, but (E) hinder lateral spreading, as illustrated by this photograph of a water droplet placed on a small peristome sample (here, flow along the channels has been restricted experimentally). (F) The large apparent contact angle, 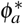, preceding lateral spreading, arises due to contact line pinning, and is determined by a combination of the maximum slope of the channel ridge, Φ, and wettability of the surface, *ϕ_a_*(*p*).

As with many plant surfaces with unusual wetting properties, the peristome is covered by a highly regular, hierarchical microstructure. This microstructure typically consists of two length scales of radially oriented channels, referred to as ‘macroscopic’ and ‘microscopic’ channels in the following (see Fig. 1 B). Each microscopic channel is formed by a single row of overlapping epidermal cells, which form a series of steps [21, 22]. By contrast, the macroscopic channels are multicellular structures visible to the naked eye, each containing multiple smaller channels. What is the function of the two different channel sizes in the context of prey capture?

The difficulty in answering this question lies in the need to disentangle the influence of the hierarchical surface topography and the intrinsic surface chemistry on (i) the ability of insects to attach to the peristome, and (ii) the peristome’s wetting properties [5]. We devised a set of experiments which enabled us to investigate each of these factors independently: The effect of the hierarchical topography was assessed by measuring friction forces of stick insect (*Carausius morosus*) adhesive pads on four surfaces, each in a wet or dry condition: (i) accurate epoxy replicas of *N. veitchii* peristomes; (ii) epoxy surfaces with rectangular channels produced by photolithography and comparable in dimensions to those of either macroscopic or microscopic peristome channels; and (iii) smooth epoxy surfaces (see Fig. S 1). The effect of surface chemistry was quantified by conducting these measurements on the same set of surfaces but with variation in their wettability. Lastly, we estimated the intrinsic wettability of the peristome through dynamic wetting measurements comparing fresh peristome samples with accurate replicas of varying wettability. In combination, these experiments allow us to separately assess the role of intrinsic wettability of the peristome cuticle and the hierarchical surface structure in the spreading and stabilisation of water films, and the slipperiness of the peristome to insect visitors, enabling us to determine the functional significance of the hierarchical ridge structure.

## Materials & Methods

### Study species and imaging

Fresh pitchers from the species *Nepenthes fusca, N. maxima, N. petiolata, N. truncta* and *N. veitchii* were collected from Kew Gardens, London, UK. Peristomes of all species were studied by light microscopy. To this end, ~0.5mm thin cross-sections were cut orthogonal to the channels with a razor blade; cross-sections were cut near the outer edge of the peristome (where channel dimensions tend to be larger), and immediately before imaging. Peristome and epoxy substrates (see below) were also studied using scanning electron microscopy (Leo Gemini 1530VP FEG-SEM). Prior to imaging, freshly cut peristome samples were coated with 5 nm of Au/Pd alloy using a Quorum Technologies K575XD sputter coater. All peristome dimensions, such as channel depth, width and period, were measured with ImageJv1.46a [23] from the light microscopy images.

### Surface production

Accurate replicas of pitcher peristomes were produced in transparent epoxy, using a soft-imprinting method [Fig. S1 A, and see 24, 25]. Peristomes were cast in silicone rubber (Polydimethylsiloxane, PDMS) in order to produce inverse moulds which were then used to cast epoxy replicas of the original peristome. Fresh peristomes were rinsed with deionized water to remove contaminants and were subsequently blow-dried with nitrogen. Uncrosslinked PDMS was produced by mixing Sylgard 184 (Dow Corning Corporation, Midland, USA) in a crosslinker:base ratio of 1:10, followed by degassing in a vacuum chamber. Cut-out pieces of the peristome were completely submerged in PDMS in order to prevent shrinking during the crosslinking process, and placed in a vacuum chamber for two minutes in order to remove interfacial air bubbles. The PDMS was then allowed to crosslink at room temperature for two days, peeled off the peristome, and cut into approximately 1×1 cm sections. A transparent, low-viscosity, low-shrinkage resin (PX672H/NC, Robnor Resins Ltd., Swindon, Wilts, UK) was mixed and placed on the PDMS peristome moulds, followed by degassing. The epoxy-covered moulds were then pressed onto 18 × 18mm glass cover slips and left to set for two days at room temperature. After curing, the PDMS moulds were carefully removed to obtain accurate and rigid peristome replicas (see Fig. S1 A).

The peristomes of all investigated species are covered with radial channels of two distinct length scales (see Fig. 1 A-C, and Tab. 1). Macroscopic channels had ridge widths ranging from 103 to 261 μm, and depths ranging from 34 to 129 μm. Microscopic channels, in turn, had widths ranging from 11 to 21 μm, and depths ranging from 3 to 7 μm (see Tab. 1. The width of the macroscopic channels increases slightly from the inside to the outside due to the radial geometry of the peristome [5]; all measurements were taken near the outer edge of the peristome, and ridge widths were measured at half-height, to achieve comparability). Substrates with channel dimensions similar to those of either macroscopic or microscopic channels were produced in epoxy using photolithography (Fig. S1B & C).

**Table 1.** Dimensions of macroscopic and microscopic channels of five *Nepenthes* species (N ≥ 3 per dimension, and n=2 per species), as well as maximum inclination angle of the macroscopic channel ridges (N ≥ 13, and n=2 per species), and critical apparent contact angle (CACA; N ≥6, and n=2 per species). All values are mean ±standard deviation.

|  | Species | Channel period | Ridge width at half-height | Channel depth | Ridge angle | CACA |
| --- | --- | --- | --- | --- | --- | --- |
| Macroscopic channels | *Nepenthes fusca* | $122 \pm 26 \mu\text{m}$ | $29 \pm 5 \mu\text{m}$ | $34 \pm 4 \mu\text{m}$ | $73 \pm 7^\circ$ | $86 \pm 6^\circ$ |
| | *Nepenthes maxima* | $103 \pm 9 \mu\text{m}$ | $24 \pm 2 \mu\text{m}$ | $27 \pm 4 \mu\text{m}$ | $71 \pm 8^\circ$ | $88 \pm 13^\circ$ |
| | *Nepenthes petiolata* | $238 \pm 26 \mu\text{m}$ | $60 \pm 2 \mu\text{m}$ | $128 \pm 10 \mu\text{m}$ | $78 \pm 5^\circ$ | $94 \pm 20^\circ$ |
| | *Nepenthes truncata* | $216 \pm 28 \mu\text{m}$ | $42 \pm 5 \mu\text{m}$ | $72 \pm 4 \mu\text{m}$ | $80 \pm 3^\circ$ | $95 \pm 18^\circ$ |
| | *Nepenthes veitchii* | $261 \pm 25 \mu\text{m}$ | $44 \pm 4 \mu\text{m}$ | $111 \pm 11 \mu\text{m}$ | $81 \pm 3^\circ$ | $99 \pm 12^\circ$ |
|  | Photolithography | 300 | 100 | 100 | – | – |
| Microscopic channels | *Nepenthes fusca* | $16 \pm 3 \mu\text{m}$ | $5 \pm 1 \mu\text{m}$ | $4 \pm 1 \mu\text{m}$ | – | – |
| | *Nepenthes maxima* | $11 \pm 1 \mu\text{m}$ | $4 \pm 1 \mu\text{m}$ | $2 \pm 1 \mu\text{m}$ | – | – |
| | *Nepenthes petiolata* | $21 \pm 3 \mu\text{m}$ | $6 \pm 1 \mu\text{m}$ | $4 \pm 1 \mu\text{m}$ | – | – |
| | *Nepenthes truncata* | $19 \pm 3 \mu\text{m}$ | $7 \pm 1 \mu\text{m}$ | $7 \pm 1 \mu\text{m}$ | – | – |
| | *Nepenthes veitchii* | $16 \pm 2 \mu\text{m}$ | $4 \pm 1 \mu\text{m}$ | $6 \pm 1 \mu\text{m}$ | – | – |
|  | Photolithography | 30 | 15 | 5 | – | – |

Silicon wafers were coated with SU-8 photoresist of the desired thickness by spin coating at 2000 rpm for 30 s (SU-8 2005 for 5μm thick films, SU-8 2100 for 100 μm thick films). After baking to dry the resist (2 min at 95 °C for SU-8 2005, 15 min at 95 °C for SU-8 2100), the films were brought into contact with a shadow mask consisting of patterns of lines of the appropriate width and spacing (see below), and subsequently exposed to UV light using a MJB4 mask aligner (SUSS MicroTec, Garching, Germany. 40 mJ cm ^−2^ for SU-8 2005, 120 mJ cm ^−2^ for SU-8 2100). The exposed regions underwent UV-triggered crosslinking and hardened, while the regions covered by the shadow mask did not, which allowed us to remove them in a subsequent development step. We produced substrates with rectangular ridges and channels similar in dimensions to the macroscopic and microscopic channels of *N. veitchii* and *N. truncata* (macroscopic channels: depth 100 μm, ridge width 100 μm, period 300 μm; microscopic channels: depth 5 μm, ridge width 15 μm, period 30 μm, see Fig. S1 B-C and Tab. 1). Before casting in PDMS (see above), the SU-8 patterns were coated with perfluorodecyltrichlorosilane (Sigma Aldrich, Poole, UK) in a vacuum chamber overnight, using approximately 100 μL of silane.

### Contact angle measurements

Dynamic contact angle measurements were carried out using a goniometer (Cam200, KSV Instruments Ltd., Helsinki, Finland), as shown schematically in Fig. 1 E. Cross sections of *Nepenthes* peristomes of approximately 3 × 10 mm in size were cut so that the channels were aligned with the short axis. The samples were placed ridge-side up on a hydrophobic surface, and the tip of the goniometer syringe was positioned approximately 1mm above a macroscopic channel. Droplets of 15 μL of deionized water were slowly expelled onto the peristome surface at a rate of 1 μL s ^−1^, using a computer-controlled stepper motor. A camera oriented parallel to the peristome channels recorded images of the water droplets at 10 Hz, and the ‘critical advancing contact angle’ (CACA) was measured as the maximum contact angle just before the water droplet spread into the adjacent macroscopic channel (see Fig. 1 E-F, as well as supplemental video V1).

In order to estimate the intrinsic contact angle of the natural peristome cuticle, we performed similar dynamic contact angle measurements on (i) epoxy peristome replicas with variable hydrophilicity, and (ii) paired smooth epoxy surfaces which underwent identical surface treatment. Epoxy replicas of peristomes were produced by cutting PDMS peristome moulds into 3 × 10 mm rectangles and placing them ridge-side-up in a petri dish. A drop of epoxy was placed on top of the mould, and cured for two days at room temperature. After curing, the epoxy was removed and, when inverted, exhibited the same surface topography as the original peristome. Comparable smooth epoxy surfaces were produced by casting epoxy against smooth PDMS moulds made from soft imprints of glass coverslips. Untreated smooth epoxy substrates were hydrophobic (static contact angle of deionized water 101 ±2°, n=10). To achieve variable hydrophilicity, we rendered surfaces hydrophilic via oxygen plasma treatment in a Femto UHP plasma cleaner (Diener electronic GmbH + Co. KG, Ebhausen, Germany), followed by varying ‘recovery’ times at ambient conditions in the laboratory. The time and power of the oxygen plasma treatment determines the density of OH-groups on the surface, and thus its wettability with water. A two-minute treatment at 100 W was the shortest time that produced almost fully wettable surfaces (i. e. static contact angles of 5 ±2°, n=12). Over the timescale of 2-5 days, the surfaces recovered much of their initial hydrophobicity [see tab. S2, and 26].

The following contact angle measurements were performed both on peristome replicas and smooth surfaces after identical recovery time (see Tab. S1): On the peristome replicas, CACA measurements were performed using the same conditions as for the natural peristomes. On smooth surfaces, dynamic contact angle measurements were performed by adding/removing a 5 μL drop to/from the surface at a rate of 0.5 μL s ^−1^; images were recorded every 80 ms. Static contact angle measurements were performed by placing a 2 μL drop on the surface; the angle was measured after the droplet was no longer moving.

### Force measurements

In order to assess the effects of surface topography, surface chemistry and the presence of water on the attachment performance of insects, we measured the friction forces generated by adhesive pads of stick insects (*Carausius morosus,* Sinety 1901). Insects were taken from a laboratory colony fed with bramble. For the measurements, the insects were immobilised by sliding them into a thin glass tube. One protruding leg was attached on its dorsal side to a piece of soldering wire mounted on the glass tube, using vinyl polysiloxane impression material (Elite HD+ light body, Zhermack, Badia Polesine, Italy). To prevent the claws from influencing the friction measurements, they were trimmed using micro-scissors [for more details, see 27, 28]. Friction forces were measured with a custom made 2D bending beam equipped with Vishay SR-4 strain gauges (Vishay Measurements Group GmbH, Heilbronn, Germany), mounted on a 3D motor positioning stage (M-126PD, Physik Instrumente, Karlsruhe, Germany). The epoxy test substrates were mounted on a glass coverlip attached to the end of the force trandsducer. Pads were brought in contact with a pre-load of 1mN for 5 s, followed by a 40 s slide at 0.05mm s^−1^ speed in a direction corresponding to a pull of the leg towards the body [27, 28]. During the slide, the normal force was kept constant using a feedback mechanism implemented in the LabVIEW control software. Peak friction forces were measured under dry and wet conditions and on substrates of different wettability: (i) untreated (hydrophobic), (ii) immediately after 2 min of oxygen plasma treatment (hydrophilic) and (iii) 2.5-3.5 h after the oxygen plasma treatment (moderately hydrophilic; comparable to a surface with advancing contact angles in the range estimated for an hypothetically smooth peristome surface. See Tab. S1 and results section).

All measurements were conducted in a randomised order under ambient conditions, and always at fresh positions on the test surfaces. Immediately before ‘wet’ measurements, a deionized water droplet of around 50 μL was placed on the substrates using a micropipette. Initial tests suggested that forces were insensitive to the amount of water placed on the surfaces. Visual inspection confirmed that the pads were fully sourrounded by water during ‘wet’ measurements.

### Statistics

Data in the text are given as mean ±standard deviation unless otherwise indicated, boxplots show the median and the 25 and 75 % quartiles, whiskers indicate 1.5 × the interquartile range. The effect of surface chemistry and topography on friction was analysed with repeated measures ANOVAS, and post-hoc analyses were conducted with paired t-tests. Effects were considered significant if *p* < 0.05. All statistical analyses were performed using R v3.4.4.

## Results and Discussion

### Macroscopic channels cause anisotropic spreading of water

Water droplets placed on fresh peristome samples rapidly spread along the macro- and microscopic channels (see video S1). This ‘wicking’ effect arises because the characteristic dimensions of the channels are well below the capillary length of water (≈2.7 mm), so that surface tension dominates gravity. In this scenario, a water droplet will spontaneously invade a channel if the progression of the triple contact line results in a reduction of the free energy in the system (see Fig. 1 D). Under the simplifying assumptions that the cross-section of the channels can be approximated as rectangular, and that the water-air interface is flat, this condition is met as long as [29, 30]:

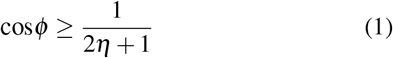

where *ϕ* is the contact angle of the wetting liquid, and *η* is the aspect ratio of the channel (depth/width). The macroscopic channels of the five *Nepenthes* species we studied satisfy η>0.3 (see Tab. 1), so that water will spread along the channels as long as the peristome is moderately hydrophilic, *ϕ_W_* < 51°.

We also studied the conditions under which water spreads *across* macroscopic channels. To this end, we cut peristome samples at right angle to the channels into thin strips, so that the triple contact line became pinned when it reached the end of the channels (Fig. 1 E & F and video S1). As a result, the water level rose above the height of the macroscopic channels. However, instead of flowing into the adjacent channel, the contact line initially remained pinned close to the top of the macroscopic ridges. A further increase in the amount of water in the wetted channel resulted in a continuous increase of the ‘apparent’ contact angle, until a critical angle was reached, at which the contact line suddenly ‘jumped’ to the next macroscopic ridge (Fig. 1 E-F, video S1). This critical apparent contact angle (CACA), was *ϕ_a_* = 93±5°(averaged across five species, see Tab. 1), significantly higher than the upper limit for spontaneous invasion along the channels. Clearly, while water spreads rapidly *along* the channels, the channel ridges present an effective barrier against the spreading of water *across* the channels [see also 4, 31, 32].

In order to quantitatively understand the lateral constraining effect of the macroscopic channel ridges, we introduce the concept of ‘Gibb’s pinning’, which occurs when an advancing three-phase contact line meets an edge-like defect [33]. Gibb’s pinning gives rise to a macroscopic ‘apparent’ contact angle 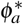 when viewed in the direction along the channel (Fig. 1 E), the magnitude of which is determined by both the geometry of the defect, and the intrinsic wettability of the surface [Fig. 1 F; The contact angle of a droplet viewed perpendicular to the channels is close to zero, an anisotropy caused by pinning effects, see 34–36]:

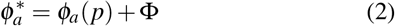

For pitcher plant peristomes, *ϕ_a_*(*p*) represents the advancing contact angle on a hypothetical peristome without macroscopic channels, and **Φ** is the maximum slope of the macroscopic channel ridges. As predicted by eq. 2, the measured CACAs increased approximately linearly with the maximum ridge slope **Φ**, exceeding it by an approximately constant amount, *ϕ_a_*(*p*) =13-18°(see Tab. 1). Hence, *ϕ_a_*(*p*) is small enough to satisfy the condition for spreading along the channels, but is also considerably larger than the expectation for a fully wettable surface (*ϕ_a_*(*p*) = 0), so increasing the barrier against lateral spreading. The combination of a moderately small value of *ϕ_a_*(*p*), and a large macroscopic ridge angle hence results in pronounced ‘wetting anisotropy’. We argue that this anisotropy serves a biological purpose: Spreading of water along the channels is crucial for successful prey capture, because it results in continuous slippery tracks running into the pitcher, so preventing sliding insects from re-gaining foothold. Lateral spreading, in turn, may be less important, as the width of the macroscopic channels is of the order of 100 μm (see Tab. 1), comparable to the width of adhesive pads of typical prey such as ants [37–41]. Lateral spreading across macroscopic channels may even be counter-productive if water is scarce, because single small droplets would no longer be able to wet the peristome continuously from the inside to the outside.

While these simple experiments suggest a clear functional role for the surface channels in terms of the directional spreading of water, they leave open if the channels influence friction forces generated by insect pads, so playing a more direct role in prey capture. In order to address these questions, we conducted friction force measurements with single adhesive pads of live stick insects (*Carausius morosus*) on a set of four epoxy surfaces: (i) accurate replicas of *N. veitchii* peristomes; surfaces with rectangular channels produced by photolithography, and comparable in dimensions to those of either (ii) macroscopic or (iii) microscopic peristome channels; and (iv) smooth surfaces cast against glass (see Fig. S 1).

### Neither roughness nor water films are sufficient to render the peristome slippery

On dry surfaces, single pad friction forces decreased significantly with increasing surface roughness (repeated measures ANOVA, F_3,51_= 50.12, p <0.001, n=18. See Fig. 2 A. This statistical analysis includes measurements on three surface types distinghuished by their wettability, see methods). However, even the lowest single pad friction forces, recorded on hydrophobic peristome replicas, were similar to the insects’ average body weight (see Fig. 2 A). Hence, the roughness of the peristome appears to be insufficient to render it slippery, consistent with results for natural pitcher plant peristomes, which are only slippery when wetted by rain or condensation [17, 42]. In seeming agreement with this observation, friction forces measured on wet hydrophobic surfaces were indeed reduced by a factor of approximately 1.5 on all surfaces. However, the peak forces generated by a single pad still amounted to at least 50% of the insects’ body weight (see Fig. 2 A). Thus, and perhaps surprisingly, neither roughness nor wetness nor their combination are sufficient conditions for peristome slipperiness.

**Figure 2.**
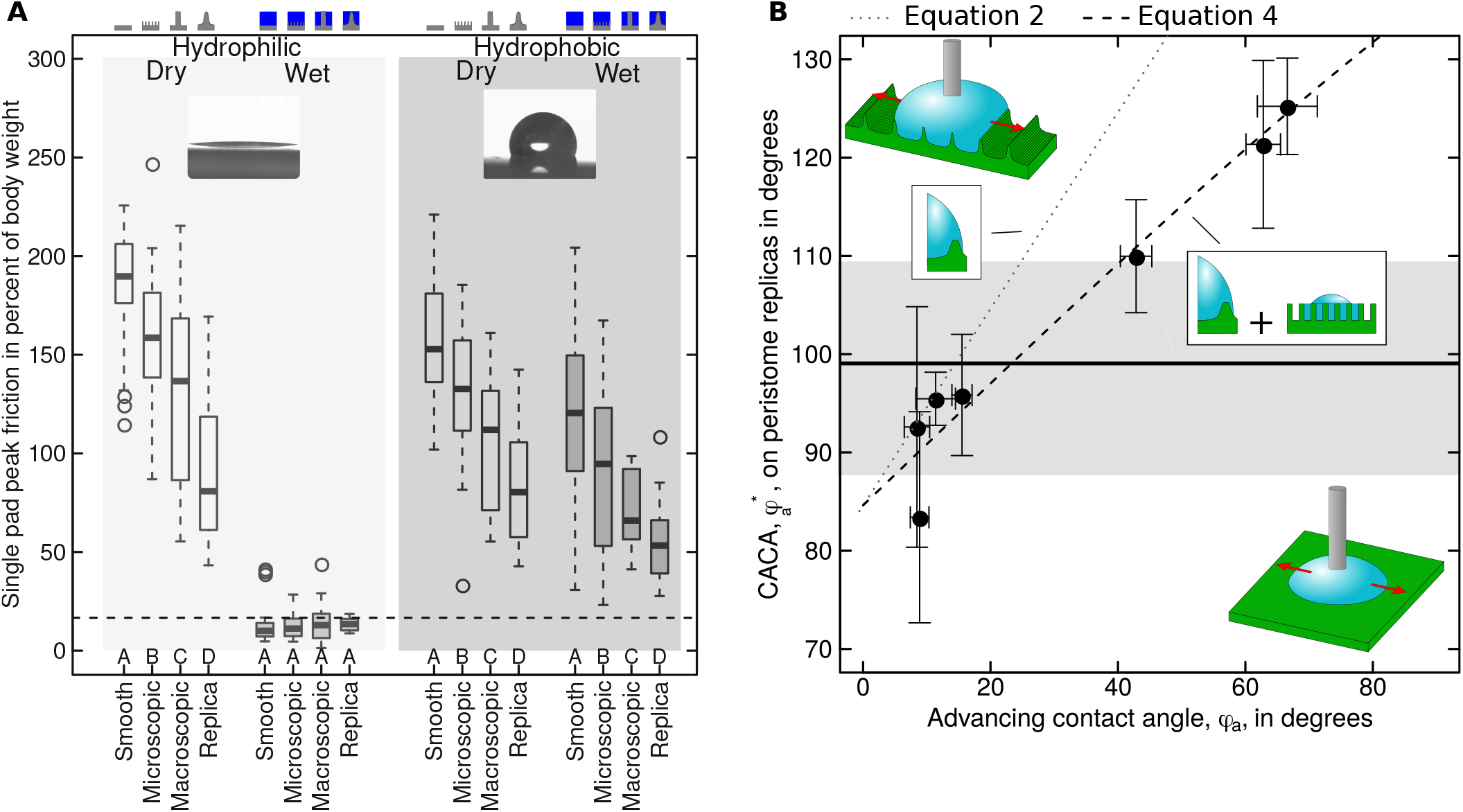
(A) In order to determine how the hierarchical structure of the peristome influences attachment performance of insects, we measured single pad friction forces of live stick insects (*Carausius morosus*) on four surfaces with controlled surface morphology (see methods). Single pad friction forces decreased with roughness and were comparable to body mass if surfaces were dry. On wet surfaces, this trend persisted if surfaces were hydrophobic, but forces dropped significantly and were no longer influenced by topography if surfaces were strongly hydrophilic (contact angles of water ≤5°). Hence, neither topography nor wetness are sufficient to render peristomes slippery. Letters indicate significant differences within each condition as obtained by paired t-tests; n=18 different insects on n=4 different surfaces for each condition; the dashed line indicates a sixth of the body mass, i. e. the value required for an insect to remain attached to the peristome with all six legs in surface contact. (B) Results from dynamic contact angle measurements on accurate replicas of *Nepenthes veitchii* peristomes (n=4-6), and smooth control surfaces with varying wettability (N=3-12 for n=3 different surfaces). Error bars show the mean ±standard deviation; the light grey area shows the 95% confidence interval of critical advancing contact angles (CACAs) measured on *N. veitchii* peristomes. The dotted grey line shows the prediction based on Gibb’s pinning; the dashed black line is the result of a non-linear orthogonal least-squares fit of a model combining Gibb’s pinning with pre-wetting of the microscopic channels (see main text).

### Insects ‘aquaplane’ when wet surfaces are strongly hydrophilic

In contrast to the hydrophobic epoxy (static contact angle of water 101±2°, n=10), the peristome of pitcher plants is readily wetted by water, suggesting a hydrophilic surface chemistry. In order to mimic natural pitcher plant surfaces more closely, we exposed all epoxy surfaces to oxygen plasma prior to friction measurements, resulting in a dramatic decrease of the static contact angle of water (5±1°, n=9).

Friction forces on dry hydrophobic vs. dry hydrophilic surfaces differed only by about 10% (repeated measures ANOVA, F_2,34_= 7.95, p <0.01, n=18), and the effect of roughness did not depend on surface chemistry (repeated measures ANOVA, F_6,102_= 1.5, p =0.18, n=18, see Fig. 2 A). However, results on *wet* hydrophilic surfaces differed quantitatively and qualitatively: First, peak friction forces were reduced by at least a factor of five, and averaged only about 15% of the insects’ body weight (see Fig. 2 A). Second, peak friction forces were no longer significantly affected by surface roughness (repeated measures ANOVA, F_3,51_= 0.36, p =0.78, n=18). Our results therefore demonstrate that peristome-like roughness is neither necessary nor sufficient to achieve ‘aquaplaning’, as a smooth hydrophilic surface is just as slippery (see Fig. 2 A). Why is a hydrophilic surface chemistry crucial to render the peristome slippery in the presence of water?

Peristome pitfall traps are activated by water, which is guided by macroscopic channels to form continous slippery tracks leading into the pitcher. However, successful prey capture also requires that water films between the insect adhesive pad and the peristome surface remain stable, and thereby prevent direct contact, causing insects to ‘aquaplane’ [18]. The low friction forces and the non-significant effect of roughness on wet hydrophilic surfaces indicate that water films between pad and surface were stable, whereas the similarity of the force measurement results between dry and wet hydrophobic surfaces suggests that water films became locally unstable. Clearly, the transition between stable and unstable water films must occur somewhere between these two extremes. How hydrophilic does the peristome need to be to avoid ‘dewetting’ of water in the contact zone?

### Surface wettability controls water film stability on smooth surfaces

Large friction forces between an insect pad and a wet peristome arise if water is completely removed from the contact zone. The initial hydrodynamic squeeze-out of water is likely fast, but the removal of the last few molecular layers poses conditions on the chemical nature of the surface, as it is driven by the minimisation of energy [18]: Dewetting of water and its replacement with the pad secretion implies an increase in the interfacial area between pad secretion and solid, but a decrease in interfacial area between water and solid, and pad secretion and water. Dewetting will only occur if the variation of energy associated with these changes in interfacial areas is negative:

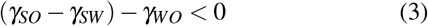

where we have assumed that the surface is smooth. Here, *γ_ij_* are the interfacial tensions between solid (*S*), water (*W*) and oily pad secretion (*O*), respectively. In the supplemental material, we show that water films will remain stable in the presence of the pad secretion as long as the contact angle of water does not exceed a critical value:

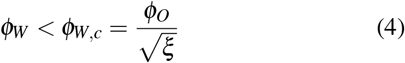

where *ϕ_i_* are the contact angles of water (*W*), and the oily pad secretion (*O*) in air, and=*γ* = *γ_w_* /*γ_o_* is the ratio of the surface tensions of the two liquids. From previous work,ξ≈2.5 [43], and *ϕ_O_* < 15°[44–46, measured on hydrophilic glass], so that water films are stable only if 0 ≤ *ϕ_W_* ≤ 10° – a remarkably narrow margin. This prediction is consistent with our results on hydrophobic and hydrophilic artificial surfaces (*ϕ_W_* = 101° vs. *ϕ_W_*= 5°, respectively), but it also raises two questions. First, our wetting experiments suggest a conservative estimate for the peristome wettability of *ϕ_a_*(*p*) > 13° – are water films unstable on natural peristomes? Second, if roughness is not required to cause insects to slip, then what is the function of the hierarchical channels in the context of prey capture? In order to address both questions, we estimated the intrinsic wettability of the peristome cuticle, and then repeated single pad friction force measurements on the set of four surfaces treated to have a comparable intrinsic wettability.

### The peristome cuticle is moderately hydrophilic

Estimating the intrinsic wettability of the peristome cuticle requires to separate the effects of microscopic roughness and surface chemistry on *ϕ_a_*(*p*), which we achieved by measuring CACAs on replicas of *N. veitchii* peristomes with varying intrinsic wettability. We varied the intrinsic wettability of these replicas by combining oxygen plasma treatment with a subsequent exposure to ambient air for controlled periods of time (contact angle ‘ageing’, see methods). As predicted by eq. 2, CACAs increased significantly with the advancing contact angle, *ϕ_a_*, measured on paired smooth epoxy surfaces which had undergone identical surface treatment. However, and in contrast to the prediction of eq. 2, the slope of this relationship was significantly smaller than one (see Fig. 2 B). This discrepancy can be attributed to the microscopic topography of the peristome, which is unaccounted for in eq. 2. In the limit of small contact angles, microscopic channels will be ‘pre-wetted’ ahead of the contact line (see inset in Fig. 2 B), so that water droplets sit on a mixture of dry and wet ‘islands’ [A Wenzel-state can be excluded based on the observation that the slope of cos 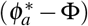 against cos*ϕ_a_* is smaller than unity. See refs. 47, 48]. The apparent contact angle *ϕ_a_* (*p*) can then be predicted as a weighted average of the respective contact angles on dry and wet patches, based on an analogy to a Cassie-Baxter model [47]:

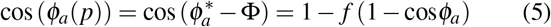

Here, *f* denotes the fraction of the pre-wetted solid surface that remains dry. A non-linear, orthogonal least-squares fit predicts *f* = 0.39 and *Φ_p_*= 85°(95% CI (0.24, 0.59) and (79°, 90°), respectively), in excellent agreement with the experimental value of **Φ**_*e*_=81 ±3°(see Fig. 2 B). Equation 5 can be used to estimate the intrinsic advancing contact angle of water on a hypothetically smooth *N. veitchii* peristome cuticle as *ϕ_a_* =25° (95% CI (13°, 44°), using the measured values for *ϕ_a_*(*p*), and the mean estimate and confidence intervals for *f* and **Φ**, respec-tively). Hence, the peristome cuticle is hydrophilic, but not fully wettable. How do insects perform on wet surfaces with comparable wettability?

### Microscopic channels enhance the stability of water films

Peak friction forces measured on wet surfaces with a moderate advancing contact angle of approximately 16±9°, within the range estimated for natural peristomes, were still significantly lower than on wet hydrophobic surfaces (see Fig. 3 A). However, and in contrast to measurements on wet hydrophilic surfaces, roughness had a significant effect (repeated measures ANOVA, F_3,51_= 8.84, p <0.001, n=18): surfaces with microscopic channels were still as slippery as in the hydrophilic condition, whereas peak friction forces on surfaces without microscopic channels were significantly larger, with a difference in means across the pooled groups of 1.36 standard deviations (Cohen’s D, 95% CI (0.64, 2.09)).

Notably, the friction performance on surfaces with or without microscopic channels differed not only quantitatively, but also qualitatively: force traces obtained on surfaces with microscopic channels were smooth and exhibited an approximately constant plateau indicative of steady-state ‘aquaplaning’. In contrast, friction forces on surfaces without microscopic channels varied considerably throughout the slide (see Fig. 3 B). These ‘stick-slip’-like fluctuations likely indicate temporary dewetting, consistent with the approximate stability margin for water films on smooth surfaces, 0 < *ϕ_W_* < 10°(see above). Two aspects of these results warrant further explanation. First, roughness appears to delay the transition from stable to unstable water films, i. e. it widens the stability margin compared to smooth surfaces. Second, this effect appears to be limited to surfaces which possess topographic features below some critical length scale.

**Figure 3.**
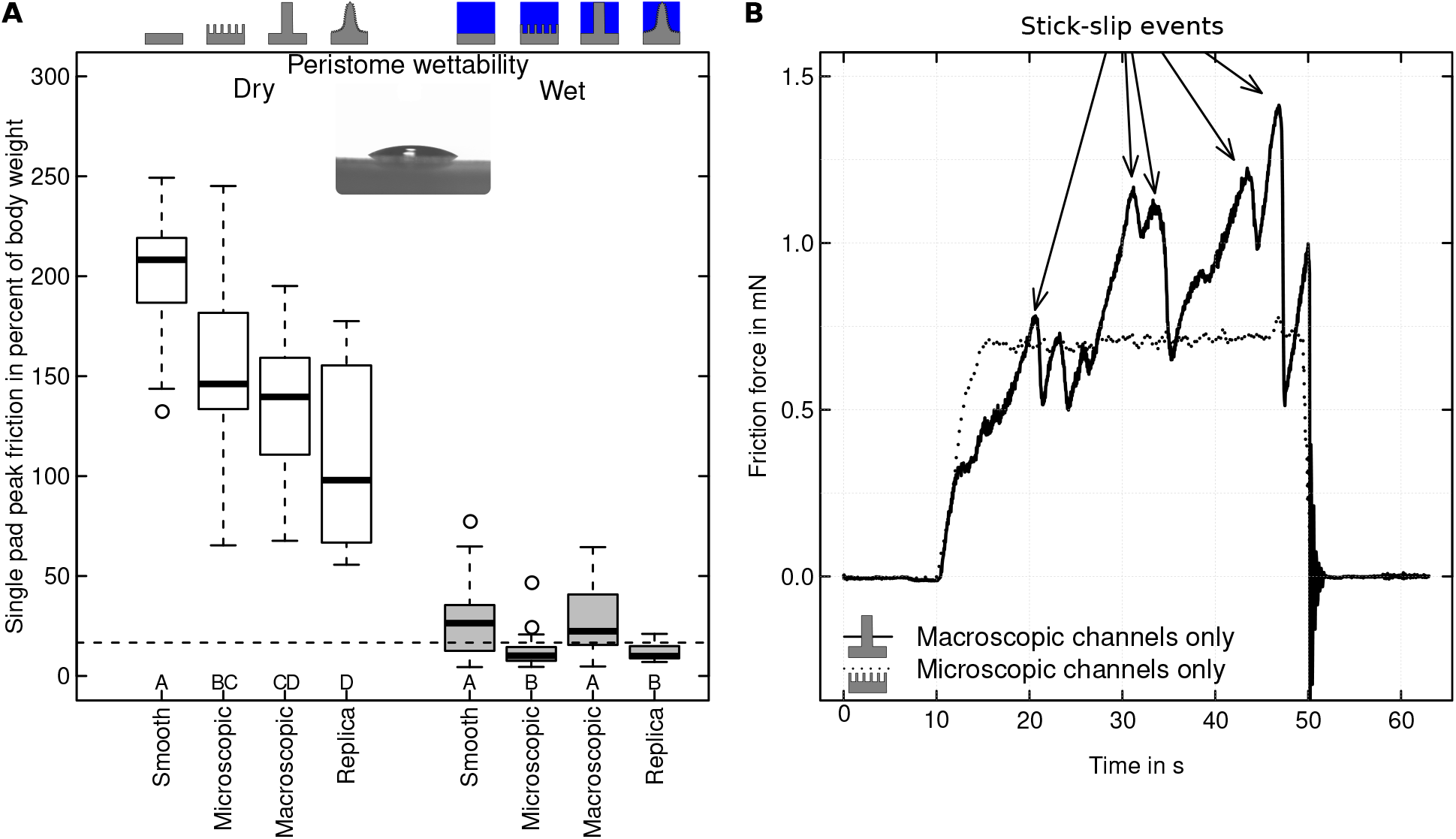
(A) Friction forces of single stick insect adhesive pads measured on surfaces with a wettability comparable to that of peristomes (advancing contact angle of 16 ±2°; *n* = 18 insects on *n* = 4 surfaces). On dry surfaces, friction forces decreased with increasing roughness, consistent with results on hydrophilic and hydrophobic surfaces. On wet surfaces, however, friction forces fell into one of two significantly different categories: peak friction was lower on surfaces with microscopic channels. Hence, microscopic channels appear to enhance the stability of water films, causing insects to aquaplane even if the surface as such is not extremely wettable. Letters indicate significant differences within each condition as obtained by paired t-tests. (B) Representative force traces of insect pads sliding on macroscopic and microscopic channel surfaces with a wettability comparable to that of the peristome cuticle. While the pads slid at a constant velocity on substrates with microscopic channels, stick-slip like instabilities occurred when microscopic channels were absent (arrows). These instabilities likely result from local dewetting of the water film, so enabling direct contact between adhesive pad and surface.

The impact of roughness on the stability of water films in the presence of the pad secretion can be assessed in analogy to the energy argument for smooth surfaces presented above. Roughness affects the interfacial area between the wetting fluid and the solid in proportion to the ‘real’ (conformal) area of contact, but the interfacial area between the two fluid phases changes only with some fraction of this area (unless water films are of molecular thickness everywhere on the rough surface). A simple assumption is that water covers the rough solid completetly, so that the water-oil interface is flat, and occupies an area equal to the the projected area of contact [for a similar argument, see 49]. In analogy to eq. 3, complete dewetting then requires:

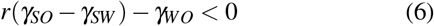

where *r* ≥ 1, is a ‘roughness factor’, defined as the ratio of the real and projected area of contact. Whether roughness widens the stability margin therefore solely depends on the sign of the bracketed term. We show in the SI that a sufficient condition for an increase in the stability margin is *ϕ_W_* < acos (*ζ*^−1^) ≈66°. Whether this increase is sufficient to keep the water films stable depends on the critical condition (a detailed derivation is provided in the SI):

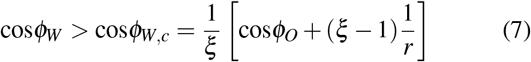

Equation 7 reduces to 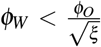 for *r* → 1 (the limit of a smooth surface). For very rough surfaces, *r* → ∞, and we find 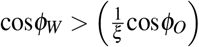, which implies that the sufficient condition for an increase of the stability margin becomes a sufficient condition for water film stability. In between these two extremes, a sufficient (conservative) stability criterion can be found by setting *ϕ_O_* = 0 in eq. 7:

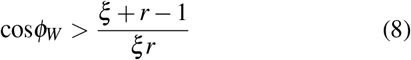

Strikingly, roughness can therefore stabilise a water film even against a completely wetting liquid. For surfaces with fractal roughness, *r* is not trivial to evaluate, but for channels with a rectangular cross-section, *r* is a simple function of channel depth and period, *d* and **p**, respectively, 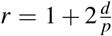. For our artificial microscopic channels, *r* = 4/3, whereas the average value for the microscopic channels of the five investigated pitcher species is *r* ≈ 9/5 (95% CI (1.46, 2.14), calculated from depth and period and assuming a rectangular cross-section), corresponding to critical values of *ϕ_W,c_* = 32 and 43°, respectively.

These approximate conditions are consistent with our experimental results, and provide physical insight into how the topography provided by the microsopic channels effectively delays the transition between stable and unstable water films, so ensuring that the peristome is slippery although the cuticle is not fully wettable.

While this physical interpretation explains the significant difference between surfaces with microscopic channels and the smooth control surface, it also implies that films should remain stable on surfaces with macroscopic channels, for which *r* = 5/3 (artificial channels), and *r*≈1.7 (peristomes, 95% CI (1.45, 2.02)), and so *ϕ_W,c_* = 41 and 42°, respectively. We suggest two possible explanations for the higher friction on surfaces with macroscopic channels. First, our definition of *r* in eq. 6 as the ratio between real and projected contact area may be invalid for macroscopic roughness with large wavelengths, as the pressure applied by the pads likely displaces some of the water. In this scenario, the interfacial area between pad secretion and water is larger than the projected contact area, so reducing the ‘effective’ roughness factor *r*. As a rough approximation, if the pads penetrated approximately 3/4 of the macroscopic channel depth, *r*≈7/6, and the conservative stability criterion yields *ϕ_Wc_* = 24°, suggesting that dewetting becomes possible. Displacement of water from within microscopic channels, in turn, is likely more difficult due to larger required pressure. Second, eq. 6 is valid if the lateral period of the roughness is small compared to the pad width, or in other words: the surface must be rough on the scale of the adhesive pad. The period of the artificial macrosopic channels is comparable to the width of stick insect (*C. morosus*) adhesive pads, *p*/*L*≈3/5, but the period of the artificial microscopic channels is an order of magnitude smaller, *p/L* ≈ 3/50.

Based on the above observations, we argue that the roughness provided by the microscopic channels is crucial to maintain the stability of water films in the contact zone. Biologically, relying on surface roughness instead of surface chemistry to ensure trapping efficacy may be advantageous for at least two reasons. First, all terrestrial plants must seal their aerial surfaces against evaporative water loss using a wax layer which consists of long-chain aliphatic hydrocarbons which are hydrophobic, so likely posing a strict limit on what can be achieved with surface chemistry alone. Second, even if these wax layers could be rendered less hydrophobic, the high surface energy required to achieve full wetting based on chemistry alone would likely attract contaminating particles and chemicals. Particle contamination strongly reduces the trapping efficiency of the peristome [50], and, as we demonstrated above, roughness can relax the conditions posed on surface chemistry considerably, in turn reducing the propensity for contamination. Combining a cuticle with moderate wettability with microscopic roughness may hence serve to maintain the peristome functional over prolonged periods of time (to the best of our knowledge, there is no evidence that Nepenthes pitcher plants make use of surfactants to wet the peristome [17], as reported for saponines in *Ruellia devosiana* [16]).

### The pitcher peristome: A multi-scale architecture to satisfy different functional demands

Pitcher plants trap insects by means of passive, water-activated pitfall traps, which requires two conditions be met:

i. Water films have to be continuous in the trapping direction, i. e. along the channels leading into the pitcher, in order to stop sliding insects from regaining foothold;
ii. Water films have to be stable against dewetting, in order to prevent direct contact between attachment pads and the peristome, thereby causing insects to ‘aquaplane’.

Our results reveal that these functional demands are satisfied by means of two distinct morphological features, separated by their characteristic length scale: Macroscopic channels restrict lateral spreading of water, and instead direct water along the radial direction (inward and outward), so creating continuous slippery tracks wider than the adhesive pads of typical prey. The stability of water films in the contact zone within these tracks, in turn, places strict demands on the surface chemistry. While the peristome cuticle is only moderately hydrophilic, it remains fully wettable and slippery due to the roughness provided by the microscopic channels, which increase the stability of water films under the adhesive pads, so causing insects to aquaplane. Together, these two mechanisms result in an efficient trapping mechanism that enables pitcher plants to capture some of nature’s most proficient climbers. Further work is necessary to understand the effect of feature-size on film stability and dewetting in more detail. Such work will suggest potential directions for the improvement of current liquid-holding surfaces inspired by the pitcher plant peristome.

## References

[1] Barthlott W, Schimmel T, Wiersch S, Koch K, Brede M, Barczewski M, Walheim S, Weis A, Kaltenmaier A, Leder A, Bohn HF. 2010 The salvinia paradox: Superhydrophobic surfaces with hydrophilic pins for air retention under water. Adv. Mater. 22: 2325–2328.

[2] Neinhuis C, Barthlott W. 1997 Characterization and distribution of water-repellent, self-cleaning plant surfaces. Ann Bot 79: 667–677.

[3] Ju J, Bai H, Zheng Y, Zhao T, Fang R, Jiang L. 2012 A multi-structural and multi-functional integrated fog collection system in cactus. Nature communications 3: 1–6.

[4] Chen H, Zhang P, Zhang L, Liu H, Jiang Y, Zhang D, Han Z, Jiang L. 2016 Continuous directional water transport on the peristome surface of *Nepenthes alata*. Nature 532: 85–89.

[5] Li J, Zheng H, Yang Z, Wang Z. 2018 Breakdown in the directional transport of droplets on the peristome of pitcher plants. Communications Physics 1: 35.

[6] Gorb S. 2009 Functional surfaces in biology: little structures with big effects., volume 1. Springer Science & Business Media.

[7] Darmanin T, Guittard F. 2015 Superhydrophobic and superoleophobic properties in nature. Materials Today 18: 273–285.

[8] Holloway PJ. 1969 Chemistry of leaf waxes in relation to wetting. J. Sci. Food Agric. 20: 124–128.

[9] Kerstiens G. 1996 Plant cuticles—an integrated functional approach. Journal of Experimental Botany 47: 50–60.

[10] Riederer M, Müller C. 2006 Biology of the Plant Cuticle. Annual Plant Reviews. Blackwell Publishing, Oxford UK.

[11] Koch K, Ensikat HJ. 2008 The hydrophobic coatings of plant surfaces: Epicuticular wax crystals and their morphologies, crystallinity and molecular self-assembly. Micron 39: 759–772.

[12] Barthlott W, Neinhuis C. 1997 Purity of the sacred lotus, or escape from contamination in biological surfaces. Planta 202: 1–8.

[13] Bhushan B, Jung YC. 2008 Wetting, adhesion and friction of superhydrophobic and hydrophilic leaves and fabricated micro/nanopatterned surfaces. Journal of Physics: Condensed Matter 20: 225010.

[14] Koch K, Bohn HF, Barthlott W. 2009 Hierarchically sculptured plant surfaces and superhydrophobicity. Langmuir 25: 14116–14120.

[15] Koch K, Barthlott W. 2009 Superhydrophobic and super-hydrophilic plant surfaces: an inspiration for biomimetic materials. Phil Trans R Soc A 367: 1487–1509.

[16] Koch K, Blecher IC, König G, Kehraus S, Barthlott W. 2009 The superhydrophilic and superoleophilic leaf surface of *Ruellia devosiana* (Acanthaceae): a biological model for spreading of water and oil on surfaces. Functional Plant Biology 36: 339–350.

[17] Bohn HF, Federle W. 2004 Insect aquaplaning: *Nepenthes* pitcher plants capture prey with the peristome, a fully wettable water-lubricated anisotropic surface. Proc Natl Acad Sci USA 101: 14138–14143.

[18] Bauer U, Federle W. 2009 The insect-trapping rim of nepenthes pitchers surface structure and function. Plant Signaling & Behavior 4: 1–5.

[19] Wong TS, Kang SH, Tang SK, Smythe EJ, Hatton BD, Grinthal A, Aizenberg J. 2011 Bioinspired self-repairing slippery surfaces with pressure-stable omniphobicity. Nature 477: 443–447.

[20] Epstein AK, Wong TS, Belisle RA, Boggs EM, Aizenberg J. 2012 Liquid-infused structured surfaces with exceptional anti-biofouling performance. Proc Natl Acad Sci USA 109: 13182.

[21] Macfarlane J. 1908 Nepenthaceae. In: Engler A, editor, Das Pflanzenreich, Wilhelm Engelmann. pp. 1–88.

[22] Heide F. 1910 Observations on the corrugated rim of *Nepenthes*. Bot. Tidsskr. 30: 133–147.

[23] Schneider C, Rasband W, Eliceiri K. 2012 NIH Image to ImageJ: 25 years of image analysis. Nat Methods 9: 671–675.

[24] Koch K, Schulte AJ, Fischer A, Gorb SN, Barthlott W. 2008 A fast, precise and low-cost replication technique for nano-and high-aspect-ratio structures of biological and artificial surfaces. Bioinspiration & Biomimetics 3: 046002–

[25] Schulte AJ, Koch K, Spaeth M, Barthlott W. 2009 Biomimetic replicas: Transfer of complex architectures with different optical properties from plant surfaces onto technical materials. Acta Biomater 5: 1848–1854.

[26] Bodas D, Khan-Malek C. 2007 Hydrophilization and hydrophobic recovery of pdms by oxygen plasma and chemical treatment—an sem investigation. Sensors and Actuators B: Chemical 123: 368–373.

[27] Drechsler P, Federle W. 2006 Biomechanics of smooth adhesive pads in insects: influence of tarsal secretion on attachment performance. J Comp Physiol A 192: 1213–1222.

[28] Labonte D, Federle W. 2013 Functionally different pads on the same foot allow control of attachment: stick insects have load-sensitive “heel” pads for friction and shear-sensitive “toe” pads for adhesion. PLoS One 8: e81943.

[29] Seemann R, Brinkmann M, Kramer EJ, Lange FF, Lipowsky R. 2005 Wetting morphologies at microstructured surfaces. Proceedings of the National Academy of Sciences 102: 1848–1852.

[30] Quéré D. 2008 Wetting and roughness. Annu Rev Mater Res 38: 71–99.

[31] Gorb S, Voigt D, Gorb E. 2007 Visualisation of small fluid droplets on biological and artificial surfaces using the cryo-sem approach. Modern research and educational topics in microscopy 2: 812–819.

[32] Box F, Thorogood C, Hui Guan J. 2019 Guided droplet transport on synthetic slippery surfaces inspired by a pitcher plant. Journal of the Royal Society Interface 16: 20190323.

[33] Oliver J, Huh C, Mason S. 1977 Resistance to spreading of liquids by sharp edges. Journal of Colloid and Interface Science 59: 568–581.

[34] Chen N, Maeda N, Tirrell M, Israelachvili J. 2005 Adhesion and friction of polymer surfaces: the effect of chain ends. Macromolecules 38: 3491–3503.

[35] Long R, Hui C. 2009 The effect of preload on the pull-off force in indentation tests of microfibre arrays. Proc R Soc A 465: 961–981.

[36] Neuhaus S, Spencer ND, Padeste C. 2012 Anisotropic wetting of microstructured surfaces as a function of surface chemistry. ACS Appl. Mater. Interfaces 4: 123–130.

[37] Moran JA. 1996 Pitcher dimorphism, prey composition and the mechanisms of prey attraction in the pitcher plant *Nepenthes Rafflesiana* in Borneo. Journal of Ecology 84: 515–525.

[38] Adam JH. 1997 Prey spectra of Bornean Nepenthes species (*Nepenthaceae*) in relation to their habitat. Pertanika Journal of Tropical Agricultural Science 20: 121–134.

[39] Bauer U, Federle W, Seidel H, Grafe TU, Ioannou CC. 2015 How to catch more prey with less effective traps: explaining the evolution of temporarily inactive traps in carnivorous pitcher plants. Proceedings of the Royal Society of London B: Biological Sciences 282: 20142675–.

[40] Labonte D, Federle W. 2015 Scaling and biomechanics of surface attachment in climbing animals. Phil Trans R Soc B 370: 20140027.

[41] Labonte D, Clemente CJ, Dittrich A, Kuo CY, Crosby AJ, Irschick DJ, Federle W. 2016 Extreme positive allometry of animal adhesive pads and the size limits of adhesion-based climbing. PNAS 113: 1297–1302.

[42] Bauer U, Bohn HF, Federle W. 2008 Harmless nectar source or deadly trap: Nepenthes pitchers are activated by rain, condensation and nectar. Proc R Soc B 275: 259–265.

[43] Hasenfuss I. 1999 The adhesive devices in larvae of *Lepidoptera* (Insecta, Pterygota). Zoomorph 119: 143–162.

[44] Federle W, Riehle M, Curtis AS, Full R. 2002 An integrative study of insect adhesion: Mechanics and wet adhesion of pretarsal pads in ants. Integr Comp Biol 42: 1100–1106.

[45] Dirks JH. 2009 Mechanisms of fluid-based adhesion in insects. Ph.D. thesis, University of Cambridge.

[46] Labonte D. 2010. Biomechanics of attachment in insects - the influence of surface energy. Bachelor Thesis; Hochschule Bremen.

[47] Bico J, Tordeux C, Quéré D. 2001 Rough wetting. EPL (Europhysics Letters) 55: 214.

[48] Bico J, Thiele U, Quéré D. 2002 Wetting of textured surfaces. Colloids and Surfaces A: Physicochemical and Engineering Aspects 206: 41–46.

[49] Kim T, Kim W. 2018 Viscous dewetting of metastable liquid films on substrates with microgrooves. Journal of colloid and interface science 520: 11–18.

[50] Thornham DG, Smith JM, Ulmar Grafe T, Federle W. 2012 Setting the trap: cleaning behaviour of *Camponotus schmitzi* ants increases long-term capture efficiency of their pitcher plant host, *Nepenthes bicalcarata*. Funct Ecol 26: 11–19.

