## Supplementary Information for "Disentangling the role of surface topography and intrinsic wettability in the prey capture mechanism of *Nepenthes* pitcher plants"

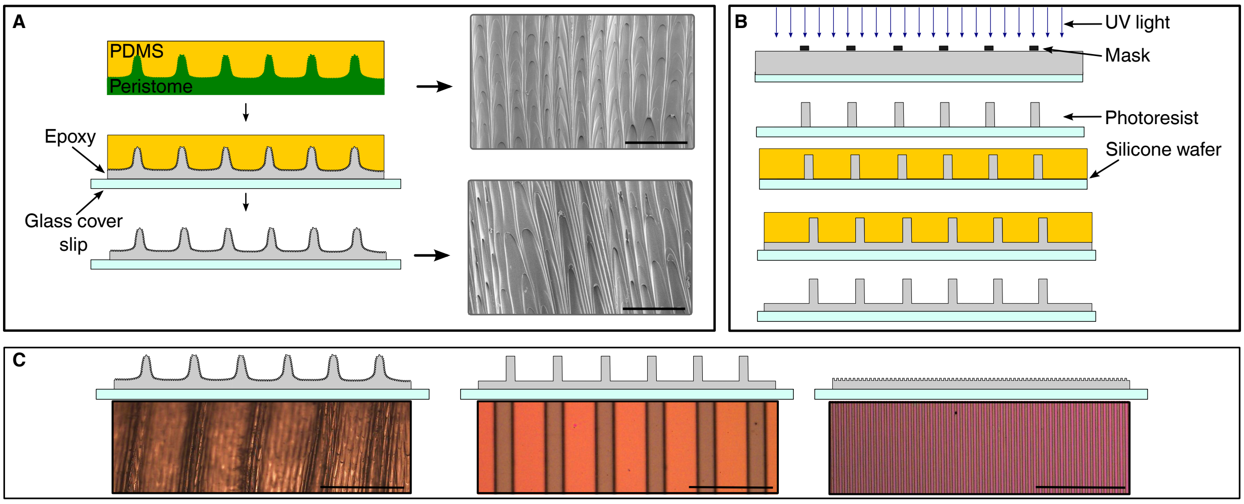


Figure S1 | (A) Replicas of peristome samples were produced by soft imprinting; the resulting Polydimethylsiloxane (PDMS) moulds were cast with epoxy, yielding accurate rigid replicas as demonstrated with scanning electron microscopy (scale bars 100 µm). (B) Surfaces with channels similar in dimension to either macroscopic or microscopic peristome channels were produced by photolithography, followed by soft imprinting with PDMS, and casting in epoxy as for the peristome replicas. (C) Light microscopy images of the produced surfaces used for force measurements (smooth reference sample not shown, scale bars 500 µm).

### Critical conditions for film stability

#### Smooth surfaces

We consider the situation of a smooth surface wetted by water, approached by an adhesive pad covered in an oily secretion. The stability of the water film can be assessed by evaluating the change in energy per unit area of contact, $A$, under the assumption that the pad secretion replaces the water. Such a replacement will increase the interfacial area between the solid and secretion, but decrease the interfacial area between secretion and water, and water and the solid, so that:

$$\frac{dE}{A} = \gamma_{SO} - (\gamma_{SW} + \gamma_{WO}) \tag{1}$$

where $\gamma_{ij}$ refers to the interfacial tensions between the oily pad secretion ($O$), water ($W$) and the surface ($S$), respectively. Dewetting will occur if $dE < 0$, and to evaluate this condition in more detail, we rewrite the above expression via the Young's equations for both oily secretion and water droplets in air (gas, $G$) on the same solid, respectively (the form of the equations requires partial wetting, but the key conclusions are unaffected irrespective of whether surfaces are fully wettable or not):

$$\gamma_{SO} = -\cos\phi_O \gamma_{OG} + \gamma_{SG} \tag{2}$$

and:

$$\gamma_{SW} = -cos\phi_W \gamma_{WG} + \gamma_{SG} \tag{3}$$

We find:


*

$$\frac{dE}{A} = (cos\phi_W \gamma_{WG} - \cos\phi_O \gamma_{OG}) - \gamma_{WO} \tag{4}$$

We will re-write this expression further in two steps. First, we make use of Good's approximation to estimate the surface tension between pad secretion and water (Good and Girifalco, 1960):

$$\gamma_{WO} = \left(\gamma_{OG} + \gamma_{WG} - 2\sqrt{\gamma^d_{OG}\gamma^d_{WG}}\right) \tag{5}$$

Here, $\gamma^d_{ij}$ are the dispersive components of the surface tension of oily pad secretion and water, respectively. For water, $\gamma^d_{WG} \approx$ 22 mN m$^{-1}$ (Fowkes, 1964), which is almost identical to available estimates for the surface tension of the pad secretion which has an oily continuous phase with negligble polar components (Hasenfuss, 1999; Dirks, 2009; Labonte, 2010), so that $2\sqrt{\gamma^d_{OG}\gamma^d_{WG}} \approxeq 2\gamma_{OG}$, and hence:

$$\gamma_{WO} = \gamma_{WG} - \gamma_{OG} \tag{6}$$

Second, we introduce a proportionality constant linking the surface tension of water and pad secretion, $\gamma_{WG} = \xi\gamma_{OG}$. Our initial energy balance now reads:

$$\frac{dE}{A} = \gamma_{OG}\left[(\xi\cos\phi_W - \cos\phi_O) - (\xi - 1)\right] \tag{7}$$

The water film is stable, if and if only the bracketed term on the right-hand side is positive, which yields:

$$\cos\phi_W > \left(1 - \frac{\phi_O^2}{2\xi}\right) \tag{8}$$

where we used the small angle approximation.[1]

This condition can only be met if $\phi_W < \phi_O$ (as $\xi > 1$), so that we can again use the small angle approximation to find:

$$\phi_W < \frac{\phi_O}{\sqrt{\xi}} \tag{9}$$

Hence, the contact angle of the pad secretion must be larger than that of water by a factor approximately equal to the square-root of the proportionality constant between the surface tensions of water and the pad secretion, $\xi \approx 2.5$, which is the result used in the main manuscript.

#### Rough surfaces

In principle, the stability of liquid films on rough surfaces can be assessed in analogy to the energy argument presented above, but a subtle difference exists: The interfacial area between the two fluid phases and the solid changes in proportion to the 'real' (conformal) area of contact (assuming complete dewetting, see below), but the interfacial area between the two fluid phases changes only with the projected area of contact (assuming a flat interfluid interface). This simple argument translates into a change of energy per unit contact area:

$$\frac{dE}{A} = r(\gamma_{SO} - \gamma_{SW}) - \gamma_{WO} \tag{10}$$

where $r \geq 1$ is defined as the area increase per unit projected area due to roughness. Hence, as long as $(\gamma_{SO} - \gamma_{SW}) > 0$, roughness will tend to increase the stability of water films in the presence of the pad secretion. Whether this widening of the stability margin is sufficient to stabilise the water films can be assessed by again using Young's equation, Good's approximation, and the proportionality constant $\xi$, in analogy to the procedure presented for smooth surfaces:

$$(\cos\phi_w\xi - \cos\phi_O) - (\xi - 1)\frac{1}{r} > 0 \tag{11}$$

It is instructive to consider two limiting cases. In the limit of shallow channels approaching a flat surface, $r \to 1$, and we recover the stability condition for smooth surfaces, i. e. $\phi_W < \frac{\phi_O}{\sqrt{\xi}}$. In the limit of deep channels with a small period, $r \to \infty$, and we find $\cos\phi_w > \left(\frac{1}{\xi}\cos\phi_O\right)$, or in other words: as long as the bracketed term in eq. 10 is positive, water films will be stable, which has $\phi_W < \mathrm{acos}\left(\zeta^{-1}\right) \approx 66°$ as a sufficient criterion.

### References

**Dirks, J.-H.** (2009). *Mechanisms of fluid-based adhesion in insects.* Ph.D. thesis, University of Cambridge.

[1]The contact angle of the oily pad secretion is likely smaller than 40° (Federle et al., 2002; Dirks, 2009).

Table S1 | Advancing and static contact angles on smooth epoxy surfaces, and critical advancing contact angles on *Nepenthes veitchii* peristome epoxy replicas were measured at defined time intervals after samples were treated with oxygen plasma. Contact angles were measured on at least three different surfaces (mean ± standard deviation (total sample size)).

| Time after treatment (h) | Advancing contact angle (°) | Static contact angle (°) | Critical advancing contact angle (°) |
|---|---|---|---|
| Untreated | - | 101 ± 2 (n=10) | 143 ± 15 (n=5) |
| 0 | 8 ± 2 (n=3) | 5 ± 1 (n=9) | 93 ± 12 (n=5) |
| 0.5 | 9 ± 1 (n=3) | 7 ± 1 (n=10) | 83 ± 11 (n=6) |
| 1.5 | 11 ± 3 (n=3) | 11 ± 1 (n=12) | 95 ± 3 (n=7) |
| 2.5 | 16 ± 2 (n=3) | 9 ± 1 (n=9) | 96 ± 6 (n=6) |
| 22 | 43 ± 2 (n=3) | 32 ± 3 (n=10) | 110 ± 6 (n=6) |
| 92 | 63 ± 3 (n=3) | 54 ± 2 (n=10) | 121 ± 9 (n=4) |
| 267 | 67 ± 5 (n=4) | 64 ± 2 (n=10) | 125 ± 5 (n=6) |

**Federle, W., Riehle, M., Curtis, A. S. and Full, R.** (2002). An integrative study of insect adhesion: Mechanics and wet adhesion of pretarsal pads in ants. *Integr Comp Biol* **42**, 1100–1106.

**Fowkes, F. M.** (1964). Attractive forces at interfaces. *Ind Eng Chem* **56**, 40–52.

**Good, R. J. and Girifalco, L.** (1960). A theory for estimation of surface and interfacial energies. iii. estimation of surface energies of solids from contact angle data. *The Journal of Physical Chemistry* **64**, 561–565.

**Hasenfuss, I.** (1999). The adhesive devices in larvae of *Lepidoptera* (Insecta, Pterygota). *Zoomorph* **119**, 143–162.

**Labonte, D.** (2010). Biomechanics of attachment in insects - the influence of surface energy. Bachelor Thesis; Hochschule Bremen.
